# “A different gut microbiome linked to inflammation found in cirrhotic patients with and without hepatocellular carcinoma”

**DOI:** 10.1101/125575

**Authors:** Federico Piñero, Martín Vazquez, Patricia Baré, Cristian Rohr, Manuel Mendizabal, Mariela Sciara, Cristina Alonso, Fabián Fay, Marcelo Silva

## Abstract

**Background:** No specific microbiome in patients with hepatocellular carcinoma (HCC) has been reported to date.

**Aim:** To compare the gut microbiome found in cirrhotic patients with and without HCC.

**Methods:** From 407 patients with Child Pugh A/B cirrhosis prospectively followed, 25 with HCC (cases) were matched with 25 without HCC (wo-HCC) in a 1:1 ratio according to age, gender, etiology, Child Pugh and severity of portal hypertension. In addition results were also compared with 25 healthy subjects. Faecal stool samples were collected noninvasively, aliquoted for DNA extraction and sequenced for the V3-V4 region of the microbial 16S rRNA (Illumina MiSeq Platform).

**Results:** There were no significant clinical differences among cases and controls. We found a differential abundance in family members of Firmicutes with a 3-fold increased of Erysipelotrichaceae and a 5-fold decrease in family Leuconostocaceae in HCC when compared to wo-HCC controls. Genus Fusobacterium was found 5-fold decreased in HCC versus wo-HCC. The ratio bacteriodes/prevotella was increased in HCC due to the significant decrease in the genus prevotella. Genus Odoribacter and Butyricimonas were more differentialy abundant in HCC. This pattern has been previously associated with an inflammatory milieu with a putative increased activation of NOD-like receptor signalling pathways. A Random Forest model trained with differential abundant taxa correctly classifyed HCC individuals with an error of 22%.

**Conclusions:** A pattern of microbiome linked to inflammation was observed in cirrhotic patients with HCC. These findings open the discussion whether or not microbiota has a physiopathologic role in HCC development in cirrhosis.

## 1. Introduction

Increasing interest has been focused during the last years regarding microbiome and human diseases including cirrhosis, alcoholic liver disease, fatty liver and fibrosis progression. Changes in gut microbiome have been observed with progression of liver disease ^1–4^ including a reduced abundance of taxa considered benign, such as Lachnospiraceae, Ruminococcaceae and Clostridialies and a higher abundance of Enterobacteriaceae and Bacteriodaceae ^2^. This gut microbiome profile remains stable unless a decompensation of cirrhosis appears with a relative increase in Enterobacteriaceae and Bacteriodaceae ^2,5^.

A renewed novel research has focused on microbiome and cancer development. However, scarce data has been written regarding microbiome and hepatocellular carcinoma (HCC). In patients with cirrhosis, there is an annual risk rate between 1 to 6% of developing HCC ^6–8^. Previous studies showed that the severity of liver disease and chronic inflamation ^9^ are predictors for development of this tumor ^10,11^. These changes have been linked to an inflammatory pro-oncogenic microenvironment from the intestine to the liver in murine models ^12^.

Inflamatory signals derived from a change in intestinal microbiome has been proposed as a novel carcinogenic mechanism ^12–14^. This gut-liver axis may also become a potentially preventive therapeutic target ^15,16^. Therefore, the aim of the present study was to compare the gut microbiome found in patients with cirrhosis with and without HCC, in order to try to identify a specific gut microbiome profile among cirrhotic patients with HCC.

## 2. Materials and Methods

### 2.1 Study design, setting and participating centers

This observational case-control study was nested on a prospective longitudinal cohort of patients with cirrhosis who were followed-up in our Liver Unit at Austral University Hospital, School of Medicine, in colaboration with HERITAS (Rosario), CONICET and the National Academy of Medicine from Argentina. This study was carried out between December 2015 and October 2016 in accordance with international recommendations for observational studies^17^.

### 2.2 Eligibility criteria

A consecutive non-probability sampling of adult subjects (>17 years old) with clinical or histological diagnosis of cirrhosis, functional status Child Pugh class A or B was included. Clinical or histological diagnosis of cirrhosis was done according to international consensus guidelines ^18^. Exclusion criteria consisted of any of the following: a) Previous liver transplant, b) patients under any immunosuppressive treatment, c) prior or current treatment with any pre- or probiotic, d) active alcoholism (cessation of alcohol intake at least 3 months prior to study entry), e) past or present history of neoplasms, f) any major surgery or severe traumatic injury within 28 days prior to inclusion, g) active infection grade >2, according to the NCI CTCAE criteria, version 4.0 ^19^, h) infection with human immunodeficient virus (HIV), i) any antibiotic treatment should have been completed one month prior to the inclusion, excluding primary or secondary prophylaxis for bacterial infections in cirrhosis, j) diarrhea secondary to any commensal, including Clostridium difficile diarrhea within 6 months prior to the inclusion of the subject in the study, k) any malabsorption disorder, celiac disease or inflammatory bowel disease including ulcerative colitis and Crohn’s disease.

Some dietary habits were precluded in all patients including high fat diets and alcohol consumption prior the inclusion to this study. All the patients lived in urban areas with similar dietary habits and water consumption.

### 2.3 Selection of cases and controls

Cases were defined as those patients meeting the above eligibility criteria and with imaging or histological diagnosis of HCC. Imaging HCC diagnosis was performed with a tri-phase dynamic study, either Computerized Axial Tomography (CT) or Magnetic Resonance Imaging (MRI) as recommended by international guidelines ^20,21^. All cases were allowed to have a history of HCC treatment, excluding liver transplantation. They should have at least one active HCC nodule ^22^ at time of fecal stool sample collection. Images were centrally reevaluated by a single observer, blinded to clinical and exposure variables.

Every case was matched with one control without HCC (wo-HCC) in a 1:1 ratio according to age, gender, etiology of liver disease, Child Pugh score and presence of clinically significant portal hypertension. Exclusion of HCC in all controls was done with either CT or MRI scans during the screening period. Selection process of cases and controls was prior to stool samples collection and was blinded from results of gut microbiome.

### 2.4 Healthy controls

In addition, we studied a gut microbiome dataset of 25 matching healthy controls. These subjects were included in a previous epidemiological microbiome research from Rosario, Argentina ^22^. This dataset was validated using the Illumina MiSeq technology as described below exclusively for this work.

### 2.5 Exposure variables: definition and meassurement

The following measurements were recorded at baseline in each subject enrolled:

. *Demographic data*: age, weight and height, concomitant medications including use of lactulose, rifaximin or norfloxacin.
. *Laboratory* including red and white blood cells count, platelets, liver function tests, urea, creatinine, plasma electrolytes, prothrombin time, INR (International Normatized Ratio) and serum alpha-fetoprotein (AFP). These blood tests were measured in the same day or the day after fecal stool sample collection in all subjects, blinded from clinical variables and microbiome results.
. *Clinical variables*: complete physical examination and past medical history, Child Pugh Score and presence or past history of clinically significant portal hypertension was evaluated during the screening period and revaluated at the same day of fecal sample stool collection. The latter included presence of ascites, hepatic encephalopathy (HE; West Haven grade criteria)^23^ and presence of esophageal or gastric varices. The degree of ascites was classified as mild, moderate or severe according to ultrasonographic or physical examination.
. *Tumor characteristics*: Subjects with HCC diagnosis were staged according to Barcelona Clinic Liver Cancer staging (BCLC) ^20,21,24^ at study entry. Type of previous tumor treatment was registered: trans-arterial chemoembolization (TACE), radiofrequency ablation (RFA), percutaneous ethanol injection (PEI) and liver resection (LR).

### 2.6 Fecal stool samples collection, DNA extraction and sequencing

Fecal samples were obtained noninvasively either at home or in the hospital, in a plastic collection kit at any time during the day. All samples were stored at −70°C. Samples obtained at patient’s home, were maintained for 24hs at 4 °C, until they were taken to the hospital and stored at −70 °C.

Each fecal sample was aliquoted for final processing in Heritas laboratory in Rosario, Argentina. Total DNA extraction from stool samples (about 200 mg) was performed using QIAmp DNA Stool Mini Kit following manufacters’s instructions. The 16S rRNA V3-V4 hypervariable region was firstly amplified using PCR method (20 cycles) and then a second reading for sample identification (6 cycles). Amplicons were cleaned using Ampure DNA capture beads (Argencourt-Beckman Coulter, Inc.) and quantified using Quanti-iTTM PicoGreen^®^ DNA Assay Kit (Invitrogen Molecular Probes, Inc., Eugene, OR, USA) with the standard protocol (high range curve – half area plate) and pooled in molar concentrations before sequencing on the Illumina MiSeq platform (Illumina, Inc., San Diego, CA, USA) using 2×300 cycles PE v3 chemistry. In each procedure 3 measurements were made to avoid information bias. The operators of this measurement were blinded to any clinical variables, including information regarding if a subject was either a case or a control.

### 2.7 Sequence data pre-processing, classification and taxonomic assignment

Amplicon sequencing produced 11639500 raw paired-end (PE) reads. We followed the 16S SOP described in Microbiome Helper ^27^. FastQC (v0.11.5) was used to analyze the raw data quality of PE reads. Paired-end reads were stitched together using PEAR (v0.9.10). Stitched reads were filtered by quality and length, using a quality score cut-off of Q30 (phred quality score) over 90% of bases and a minimum length of 400bp. Concatenated and filtered fastq sequences were converted to fasta format and we removed sequences that contain ‘N’. Potential chimeras were identified using the UCHIME algorithm, and then the chimeric sequences were removed.

The remaining sequences were clustered to operational taxonomic units (OTUs) at 97% similarity level with a open reference strategy implemented in Quantitative Insights into Microbial Ecology (QIIME; v1.91), using SortMeRNA for the reference picking against the Greengenes v13_8 97% OTU representative sequences database and SUMACLUST for *de novo* OTU picking.

### 2.8 Statistical analysis and16S rRNA bioinformatics processing

A statistical two-tailed value, α type I error or 5% (p value <0.05) was considered. Binary or dichotomous variables were expressed as frequencies or proportions and compared by Chi-square test or Fisher exact test, as appropriate. Dicrete and continuous numerical variables were expressed according to their distribution as mean (standard deviation) or medians (interquartile range) and were compared using Student t-test or Mann-Whitney U test, respectively. In case of multiple comparisons for continuous variables, a parametric ANOVA with Bonferroni correction was done. Clinical data statistical analyzes was done using STATA version 10.1.

Bioinformatics analysis of the data was done using a custom QIIME pipeline and the R package phyloseq (https://github.com/joey711/phyloseq). OTU table was rarefied at 12566 sequences per sample for alpha and beta diversity and random forest analyses. To provide alpha diversity metrics, we calculated observed species, Chao1 and Shannon’s diversity index. To evaluate beta diversity among cases, controls and healthy individuals UniFrac (weighted and unweighted) distances and Bray Curtis dissimilarity were used, prior removal of OTU’s not present in at least 5% of samples. The UniFrac and Bray Curtis measures were represented by two dimensional principal coordinates analysis (PCoA) plots. Differences between groups were tested by a permutational multivariate analysis of variance (PERMANOVA) using distance matrices function (ADONIS) implemented in the R vegan package^34^. The evaluation of beta diversity between cases and controls were also determined using a principal fitted components for dimension reduction in regression as it was used to analyze microbiome data ^25^.

To understand the differences in the composition between groups, the R package DESeq2 was used to perform differential abundance estimates. Analysis of composition of microbiomes (ANCOM) was carried out with the ANCOM R package (https://www.ncbi.nlm.nih.gov/pubmed/26028277) with 5% FDR.

We build a Random Forest model with the random Forest R package using OTU’s relative abundance as features to predict samples from HCC or wo HCC groups, measuring the “out of bag” (OOB) error to estimate the error model. From the list of variables importance, measured by the mean decrease in Gini index, we selected the three OTU’s that contributed the most to the group classification. A random forest was trained with these 3 OTU’s. To determine the model significance a permutation test with 1000 permutations was run.

### 2.9 Functional analysis

In order to evaluate the impact of any observed differences in the microbiome composition we further performed an indirect functional assessment to infer the metagenomic composition with PICRUSt (Phylogenetic Investigation of Communities by Reconstruction of Unobserved States). We removed OTU’s not present in the database used for OTU picking, the OTU table normalized by predicted 16S copy numbers was used to predict KEGG otholog (KOs) collapsed to KEGG Pathways. This PiCRUST analysis included any association with NOD-like receptor (NLR) signaling pathway^14^.

## 3. Study end-points

The objective of this study was to compare gut microbiome profile in patients with cirrhosis with and without HCC, in order to identify a specific profile of microbiome potentially associated with the risk of developing liver cancer.

### 3.1 Potential biases and confounding variables

In order to anticipate and avoid selection bias regarding HCC-cases and wo-HCC, subjects were followed and treated under the same standards of care and were selected from a cohort of patients prospectively followed at Austral University Hospital by investigators who were blinded to gut microbiome results. Acceptances and declines to participate in this study were recorded. Selection bias was avoided, excluding HCC diagnosis with a CT or MRI scan in wo-HCC controls.

### 3.2 Ethical considerations

This study protocol has been developed according to national standards for ethical, legal and regulatory requirements established in the 6677/10 ANMAT disposition, international ethical standards according the 2008 Helsinki Declaration and its amendments from the Nuremberg code, universal declaration on the human genome and human rights adopted by the General Conference of UNESCO 1997, as well as GCP standards (“Good Clinical Practice”).

## 4. Results

### 4.1 Participating Patients characteristics

From a total of 407 patients who were prospectively followed and entered in the prescreening study period, 72 patients with HCC and 190 without HCC were selected as potential cases and controls. Overall, 25 cases and 25 controls (wo-HCC) were well matched according to those variables previously described and were included in the final assessement of gut microbiome (Figure 1). A subgroup of 25 healthy individuals were also enrolled, matched according to gender in a 1:1 ratio ^26^. Baseline patient characteristics are shown on Table 1. Considering severity of liver disease, most of the patients were Child Pugh A (74%), only 10% had ascites and 8% had presence of HE.

**Figure 1.**
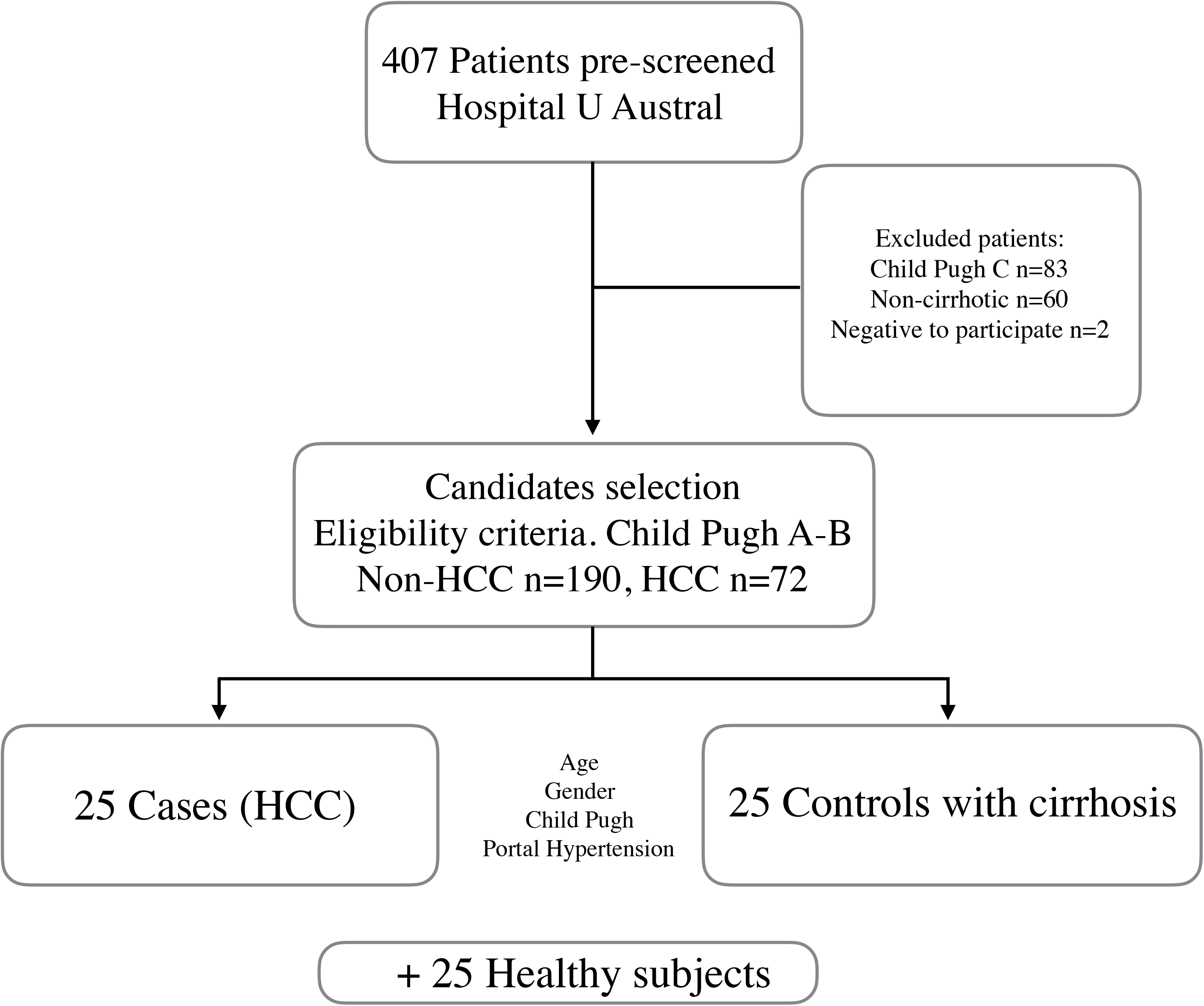
Inclusion and Exclusion Criteria flow chart.

**Table 1.** Patients’ baseline characteristics.

| **VARIABLE** | **Values** |
| --- | --- |
| **Age, years (± SD)** | **64 ± 8** |
| **Male gender, n (%)** | **44 (88.0)** |
| **BMI, Kg/m<sup>2</sup>** | **27.9 ± 4.5** |
| **Child Pugh A/B, n (%)** | **37 (74.0)/13 (26.0)** |
| **Etiology of liver disease, n (%)** |  |
| *Hepatitis C virus* | 13 (26.0) |
| *Alcohol* | 11 (22.0) |
| *NASH* | 10 (20.0) |
| *Cryptogenic* | 8 (16.0) |
| *Hepatitis B virus* | 4 (8.0) |
| *Autoimmune* | 2 (4.0) |
| *Hemochromatosis* | 2 (4.0) |
| **Ascites, n (%)** |  |
| **No** | 45 (90.0) |
| **Mild** | 2 (4.0) |
| **Moderate-severe** | 3 (6.0) |
| **Encephalopathy, n (%)** |  |
| **No** | 46 (92.0) |
| **Grade I-II** | 4 (8.0) |
| **Grade III-IV** | - |
| **Esophageal varices, n (%)** | 41 (82.0) |
| **ECOG 0-2/3-4, n (%)** | 50 (100)/ - |
| **Abbreviations:** NASH, non-alcoholic steatohepatitis. BMI, Body Mass Index. |  |

### 4.2 Cases and controls: Clinical and tumor variables

In patients with HCC the mean number of intrahepatic nodules were 2.2 ± 1.8, with a major nodule tumor diameter of 3.8 ± 1.6 cm. Considering BCLC stages, 2 patients were stage 0, 12 patients were stage A, 5 were in stage B and 6 in stage C-D. Vascular invasion was observed in 3 patients and extrahepatic spread in 4. Previous HCC treatments consisted of RFA in 1 patient, LR in 3, TACE in 17 and Sorafenib in 3 patients.

Table 2 shows baseline clinical and laboratory variables comparing cases and controls. There were no significant differences in any clinical variable. The use of rifaximin, norfloxacin or lactulose was not significantly different between cases and controls.

**Table 2.** Baseline clinical and laboratory variables among cases and controls.

| **VARIABLE** | **Cases**<br>**(HCC)**<br>**n= 25 (50%)** | **Controls**<br>**(wo-HCC)**<br>**n= 25 (50%)** | **P** |
| --- | --- | --- | --- |
| **Age, years (± SD)** | 64 ± 9 | 63 ± 7 | 0.53 |
| **Male gender, n (%)** | 22 (88.0) | 22 (88.0) | 1.0 |
| **Diabetes mellitus, n (%)** | 14 (.56.0) | 8 (32.0) | 0.17 |
| **BMI, Kg/m<sup>2</sup>** | 27.9 ± 5.4 | 28.1 ± 3.6 | 0.87 |
| **Child Pugh A/B, n (%)** | 17 (68.0)/8 (32.0) | 20 (80.0)/5 (20.0) | 0.93 |
| **MELD** | 6.7 ± 0.4 | 6.6 ± 0.5 | 0.76 |
| **Etiology of liver disease, n (%)** |  |  |  |
| *Hepatitis C virus* | 6 (24.0) | 7 (28.0) |  |
| *Alcohol* | 6 (24.0) | 5 (20.0) |  |
| *NASH* | 5 (20.0) | 5 (20.0) |  |
| *Cryptogenic* | 4 (16.0) | 4 (16.0) |  |
| *Hepatitis B virus* | 2 (8.0) | 2 (8.0) |  |
| *Autoimmune* | 1 (4.0) | 1 (4.0) |  |
| *Hemochromatosis* | 1 (4.0) | 1 (4.0) |  |
| **Ascites, n (%)** |  |  |  |
| **No** | 22 (88.0) | 23 (92.0) | 0.31 |
| **Mild** | 2 (8.0) | 0 (0.0) |  |
| **Moderate-severe** | 1 (4.0) | 2 (8.0) |  |
| **Encephalopathy, n (%)** |  |  |  |
| **No** | 23 (92.0) | 23 (92.0) | 1.0 |
| **Grade I-II** | 2 (8.0) | 2 (8.0) |  |
| **Grade III-IV** | - | - |  |
| **Portal hypertension, n (%)** | 20 (80.0) | 23 (92.0) | 0.22 |
| **History of decompensation, n (%)** | 9 (36.0) | 14 (56.0) | 0.15 |
| *Ascites* | 3 (12.0) | 6 (24) |  |
| *Variceal Hemorrhage* | 2 (8.0) | 1 (4.0) |  |
| *Encephalopathy* | 0 (0.0) | 1 (4.0) |  |
| **Rifaximin/Norfloxacin, n (%)** | 3 (12.0) | 5 (20.0) | 0.44 |
| **Lactulose, n (%)** | 2 (8.0) | 6 (24.0) | 0.12 |
| **Hemoglobin, gr/dL (± SD)** | 13.2 ± 2.4 | 13.0 ± 2.2 | 0.81 |
| **Platelets /mm<sup>3</sup> (± SD)** | 116,920 ± 57,753 | 124,640 ± 53,489 | 0.62 |
| **Creatinine, mg/dL (± SD)** | 0.88 ± 0.20 | 0.92 ± 0.28 | 0.55 |
| **Bilirrubin, mg/dL (± SD)** | 1.48 ± 0.98 | 1.26 ± 0.86 | 0.41 |
| **INR** | 1.33 ± 0.25 | 1.34 ± 0.30 | 0.93 |
| **AST**, IU/L (± SD) | 55 ± 46 | 43 ± 29 | 0.28 |
| **ALT**, IU/L (± SD) | 44 ± 32 | 38 ± 25 | 0.42 |
| **AP**, IU/L (± SD) | 150 ± 99 | 140 ± 73 | 0.68 |
| **γGT**, IU/L (± SD) | 192 ± 160 | 115 ± 91 | 0.07 |
| **AFP**, ng/mL, median (IQR) | 9.4 (3.2-218.0) | 3.2 (1.6-5.5) | 0.007 |

**Abbreviations:** AFP, alpha-fetoprotein. ALT, Alanine aminotransferase. AP, Alkaline Phosphatase. AST, Aspartate aminotransferase. γGT, Gamma glutamil transpeptidase. INR, International Normalized Ratio. NASH, non-alcoholic steatohepatitis.

### 4.3 Gut microbiome diversity of Cases, Controls and Healthy subjects

Cases and controls (wo-HCC) datasets of the gut microbiome were compared to a gut microbiome dataset of healthy subjects from the city of Rosario (Argentina). The Shannon’s alpha diversity index indicated that healthy controls showed the most diverse dataset of gut microbiome compared to cases and controls (ANOVA; *P*< 0.05) (Supplementary Figure 1; Supplementary Table 1A). The wo-HCC dataset consistently and significantly showed a less diverse gut microbiome composition than healthy subjects (Tukey’s honest significance test; *P*< 0.05; Supplementary Table 1B). There was not a significant difference in the microbiome diversity between HCC and wo-HCC controls. The HCC dataset was less diverse than healthy controls, but still more diverse than the wo-HCC dataset.

Beta diversity measures were computed with weighted and unweighted UniFrac distances and Bray Curtis dissimilarity calculated on OTUs relatives abundances within sample. Ordination was visualized using Principal Coordinates Analysis (PCoA) comparing HCC vs wo HCC, and healthy vs HCC vs wo HCC groups (Figure 2; Supplementary Figure 2A). To test statistical differences between groups ADONIS test was performed. Significant differences in the healthy vs HCC vs wo HCC comparisons were found (Supplementary Table 2B). There was not a clear differentiation between HCC and wo HCC groups in the PCoA and ADONIS results (Figure 2; Supplementary table 2A). To gain in-depth insight of cases and wo-HCC group, we performed a principal fitted components analysis with dimension reduction that showed a more precise and clear separation of the gut microbiome datasets between groups (Supplementary Figure 2B).

**Figure 2.**
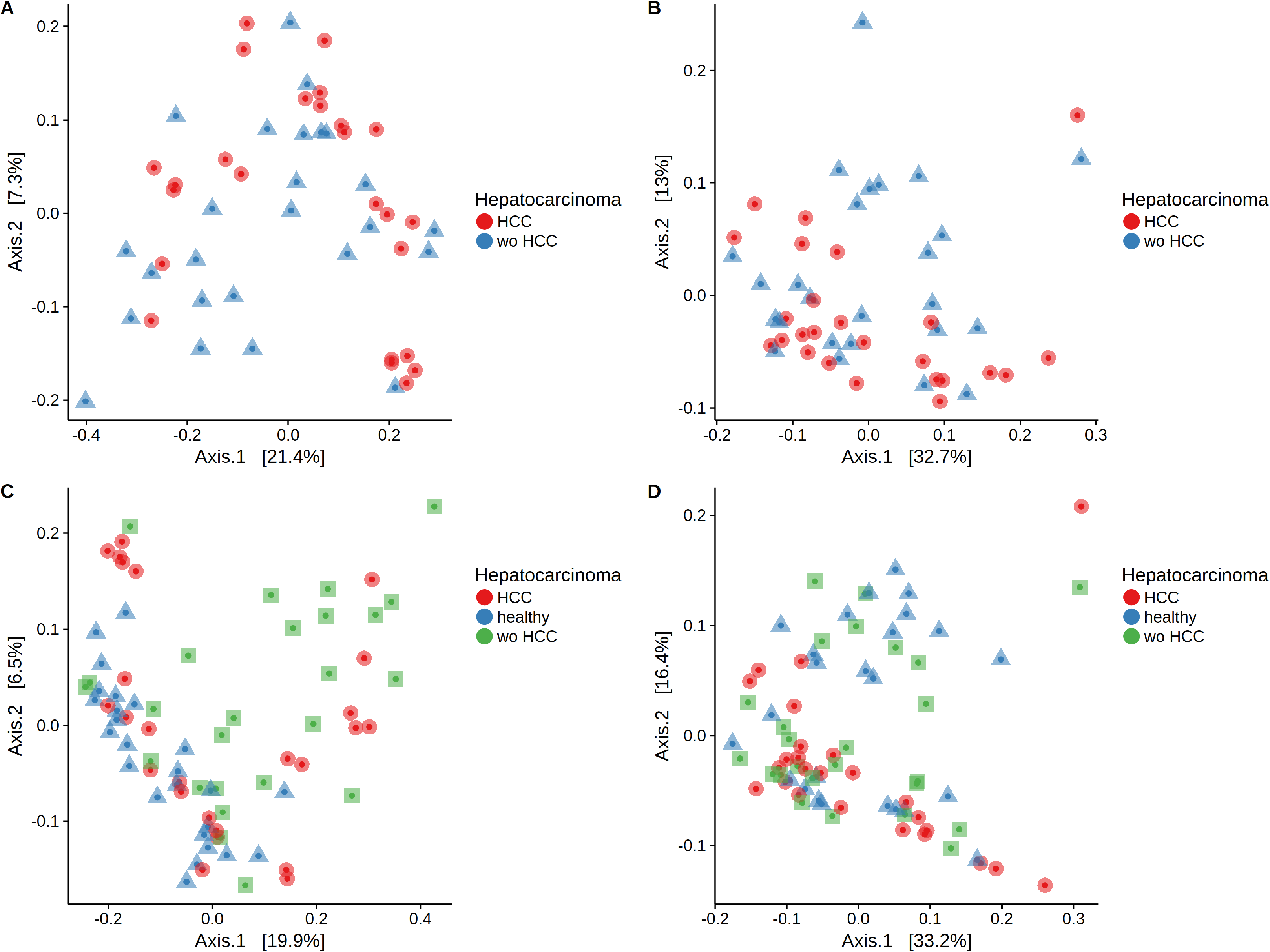
Principal Component Analysis (PCoA) plots with weighted and unweighted UniFrac distance metrics. HCC vs wo HCC a) UniFrac, b) weighted UniFrac. Healthy vs HCC vs wo HCC c) UniFrac, d) weighted UniFrac. **Note**: There was not a clear differentiation between HCC and wo HCC groups in the PCoA and ADONIS results. To gain in-depth insight of cases and wo-HCC group, we performed a principal fitted components analysis with dimension reduction that showed a more precise and clear separation of the gut microbiome datasets between groups (Supplementary Figure 2B).

### 4.4 Taxonomic distribution and differenctial abundance analysis of the gut microbiome

Taxonomic abundance at Phylum, Family and Genus levels were investigated for cases, wo-HCC and healthy groups. Differences in the relative abundance of taxa were evident at the levels of Family and Genus among the three groups. Of note is the expansion of Bacteroidaceae and reduction of Prevotellaceae in the HCC group that, in turn, is reflected at the genus level in Bacteriodes and Prevotella. Consequently, the bacteriodes/prevotella ratio is greater in HCC than in the two other groups (Figure 3A).

**Figure 3.**
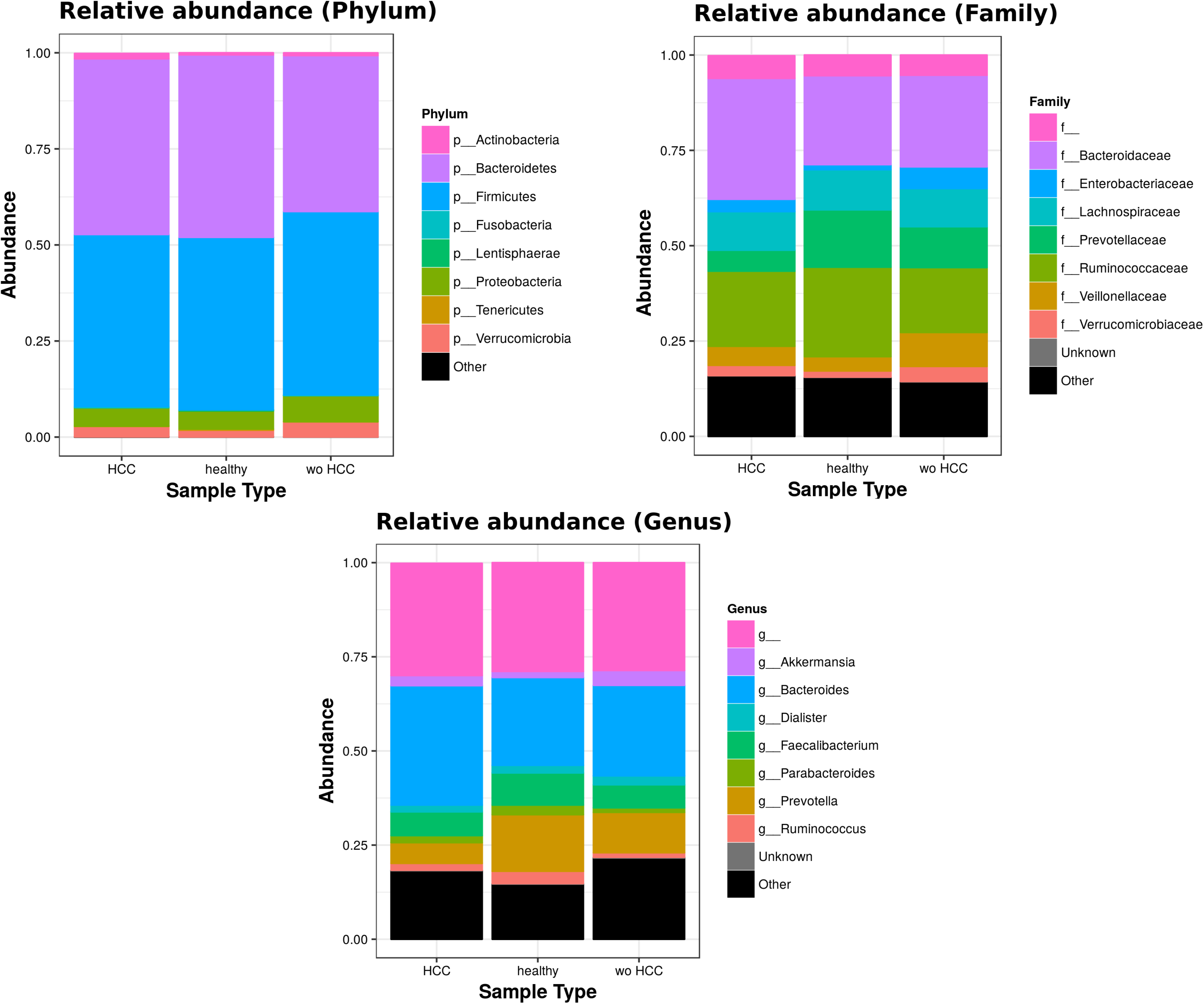
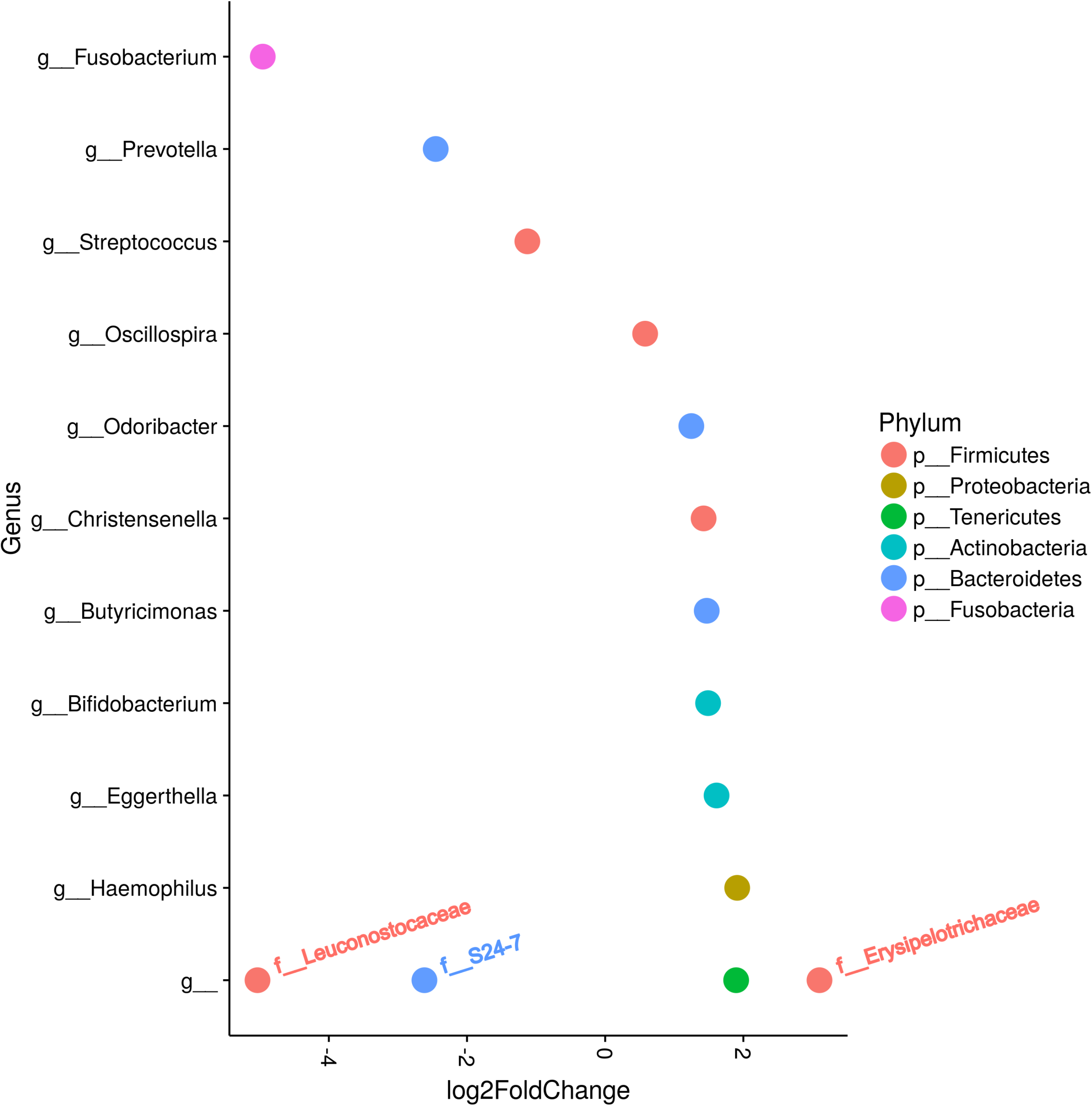
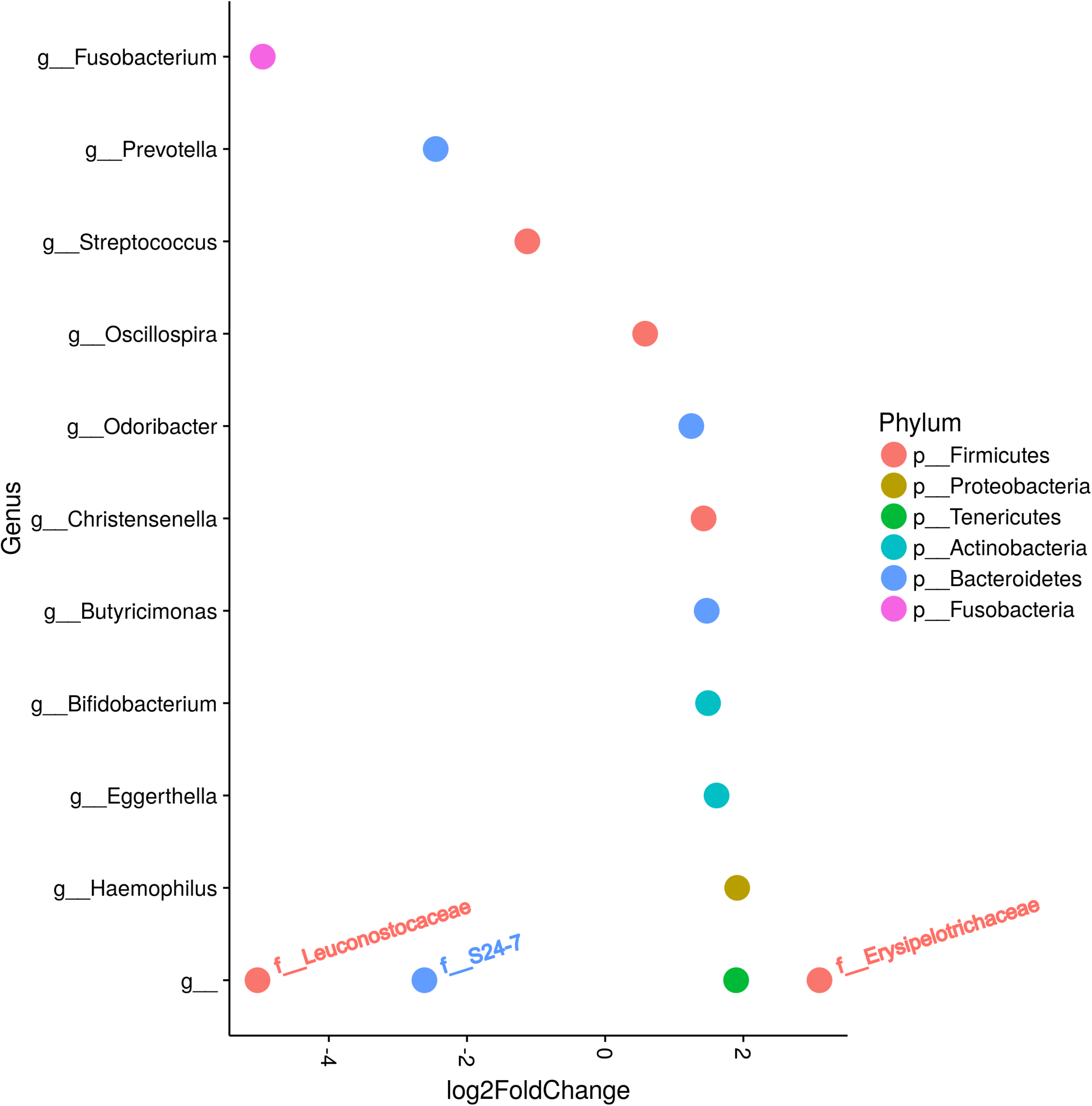
Taxonomic abundance at Phylum, Family and Genus levels between HCC, non-HCC patients and healthy individuals (Panel A). Differential abundance analysis using Phyloseq and DEseq2 (Panels B and C). **A**: A bacteriodes/prevotella ratio was greater in HCC than in non-HCC controls and healthy controls. **B**: Several genera showed interesting correlations such as Fusobacterium, Prevotella, Streptoccocus, S24-7 (Phylum Bacteroidetes) and an unknown genus (phylum Firmicutes, family Leuconostocaceae), all of which were decreased 2 to 5-fold in HCC group. The ANCOM analysis identified 3 taxa to be associated with differences between cases and controls. Two OTU’s from the Odoribacteraceae family, genus Odoribacter and Butyricimonas were also identified to be more differentialy abundant in HCC samples in the DESEQ2 analysis. The other OTU belong to the Lachnospiraceae family genus Dorea. C: Several OTUs in Prevotella genus decreased in HCC cases.

To further investigate which taxa is accounting for the observed differences in alpha and beta diversity of the gut microbiome in the HCC group, we performed a differential abundance analysis using phyloseq and DEseq2 (Figure 3B). Several genera showed interesting correlations such as Fusobacterium, Prevotella, Streptoccocus, S24-7 (Phylum Bacteroidetes) and an unknown genus (phylum Firmicutes, family Leuconostocaceae), all of which were decreased 2 to 5-fold in HCC group. On the other hand, Haemophilus, Eggerthella, Bifidobacterium, Butyricimonas, Christensella, Odoribacter, an unknown genus (phylum Tenericutes) and an unknown genus (phylum Firmicutes, family Erysipelotrichaceae) were all elevated by 2 to 3-fold in HCC group (Figure 3A-B). Each genus in Figure 3 is the result of the mean abundance of several OTUs contained in that genus and only those with significant differential abundances with an adjusted *P* value of 0.05 are shown.

Individual OTUs within some of the genera showed greater o lower values of fold change in the HCC group. For instance, several OTUs in Prevotella genus decreased between 3 to 6-fold but other Prevotella OTUs increased 1.5 to 3-fold. The mean fold change for genus Prevotella indicated a significant 3-fold decreased as shown in Figure 3A-B. A full panorama involving the complete dataset of OTUs analyzed with significant changes is observed in Figure 3C. Interestingly, all the changes in abundance in Figure 3 A-C correlated with an increase in the predicted metabolism of NLR signaling pathway in the HCC group (Figure 4). The NLR signaling pathway was previously reported to be involved in inflammatory and autoinflammatory processes ^14^.

**Figure 4.**
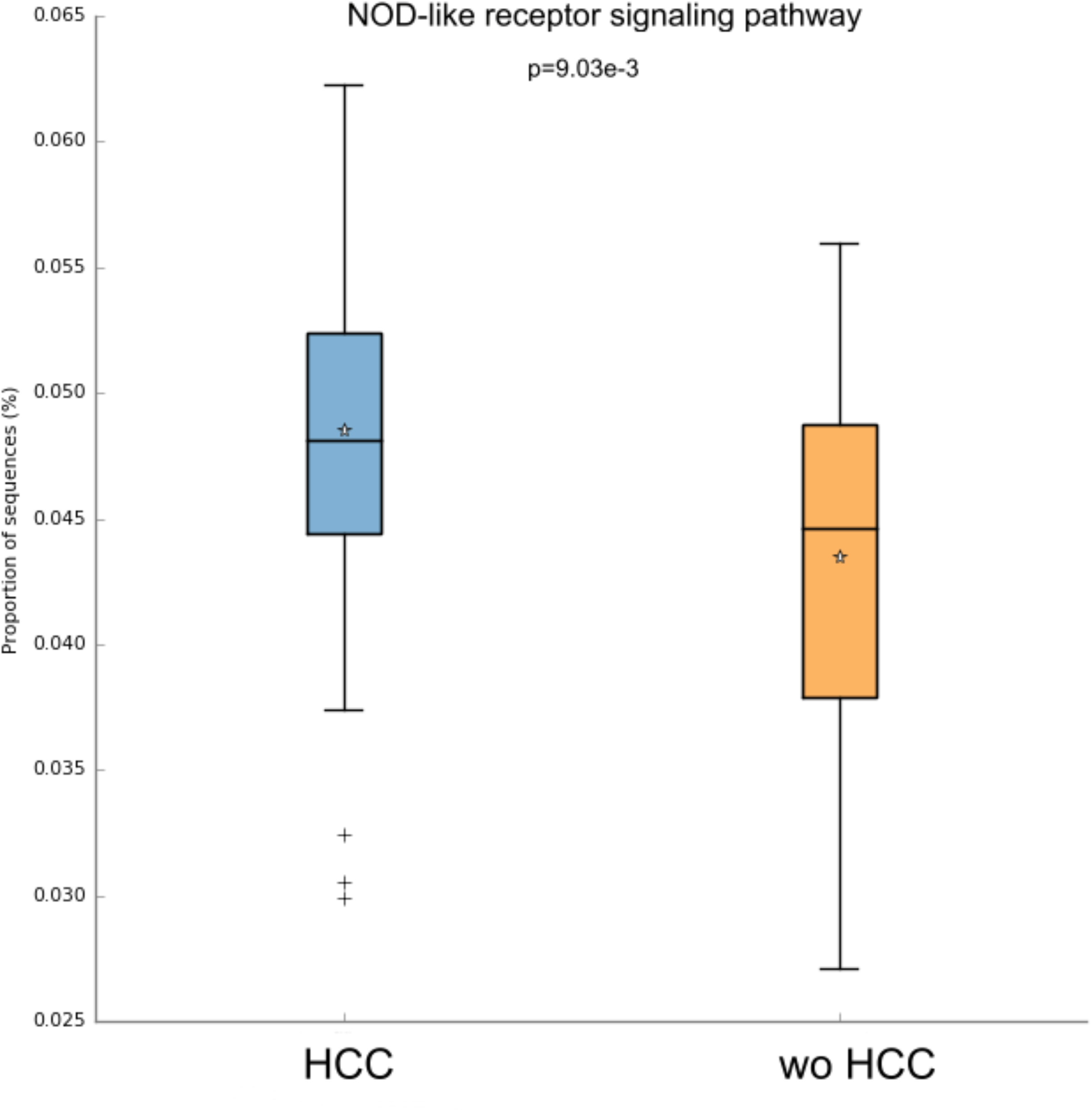
NOD-like receptor signaling pathway in the HCC and non-HCC groups. **Note**: Interestingly, all the changes in abundance in HCC patients correlated with an increase in the predicted metabolism of NOD-like receptor signaling pathway (NLR). The NLR signaling pathway was previously reported to be involved in inflammatory and autoinflammatory processes ^14^.

The ANCOM analysis identified 3 taxa associated with differences in the composition of gut microbiome between cases and controls (Figure 5A). Two OTU’s from the Odoribacteraceae family, genus Odoribacter and Butyricimonas (OTU’s ID: X4454586; X988375), were also identified to be more differentialy abundant in HCC samples in the DESEQ2 analysis. The other OTU with a decreased abundace in the HCC individuals belongs to the Lachnospiraceae family genus Dorea (OTU’s ID: X310608).

**Figure 5.**
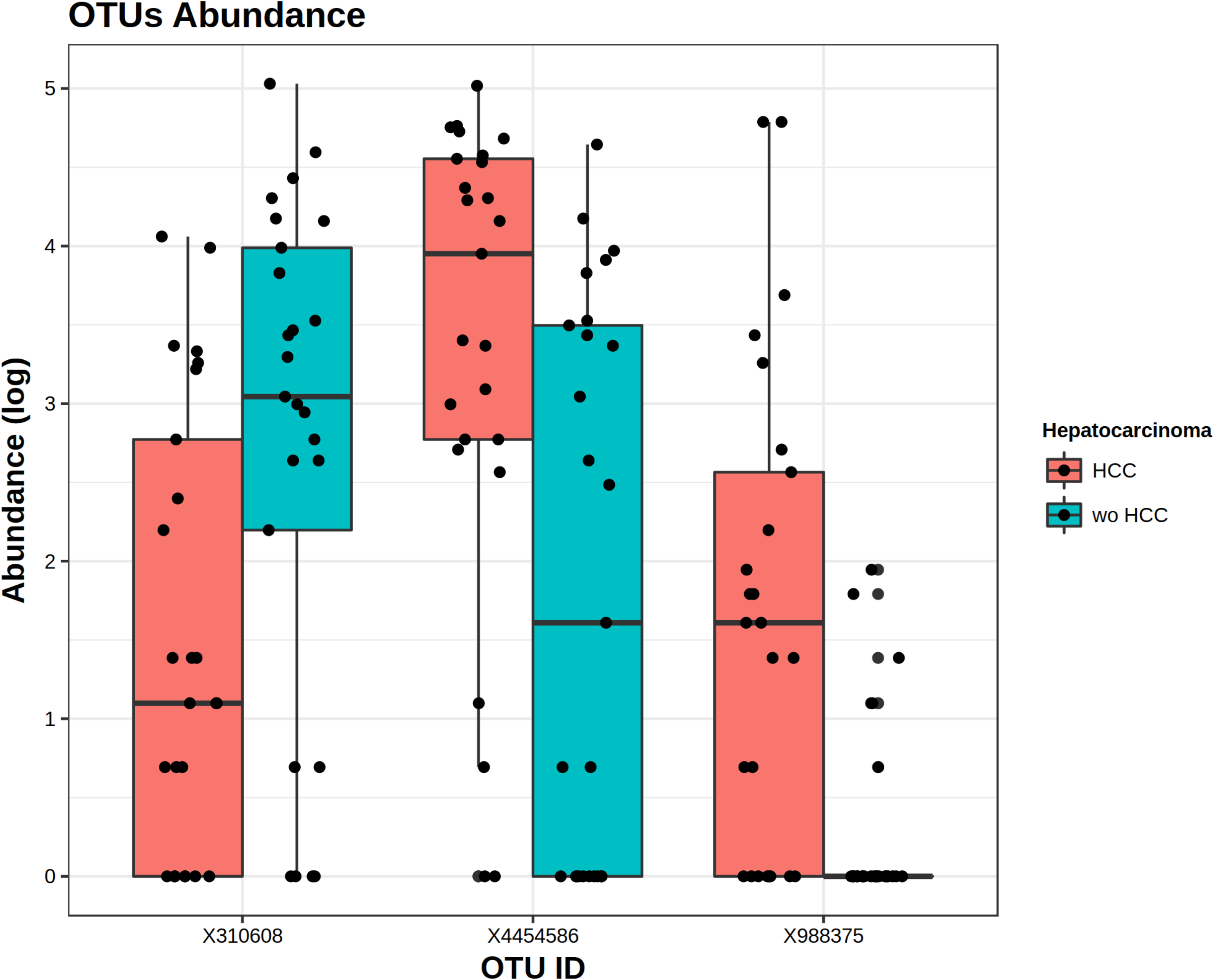
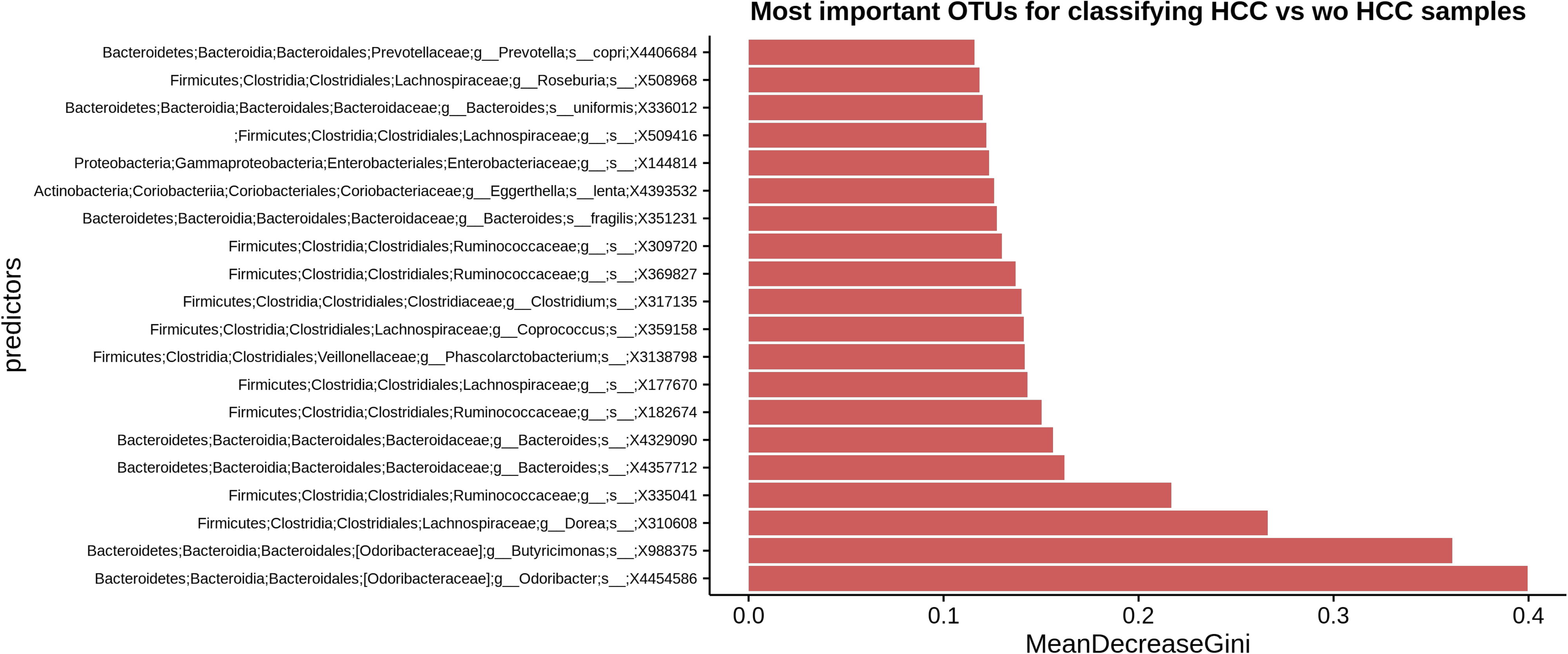
The ANCOM analysis for identifying operational taxonomic units (OTUs)(Panel A). Mean decrease in Gini index for the top ten predictor OTUs (Panel B). **A**: OTUs found in the ANCOM analysis (abundance is log transformed) identified 3 taxa to be associated with differences between cases and controls from the Odoribacteraceae family, genus Odoribacter and Butyricimonas (OTU’s ID: X4454586; X988375). The other OTU belongs to the Lachnospiraceae family genus Dorea (OTU’s ID: X310608). **B**: An analysis using OTU’s relative abundance as HCC group predictor was performed. The error OOB in the prediction was 50%, a strong indicator that it was not a good predictor of HCC or without HCC. However, observing what OTU’s were the ones that predicted better any differences in the groups, we found that the top three OTU’s were the same that in the ANCOM analysis.

### 4.5 Random Forest Analysis

We developed a Random Forests classifier using OTU’s relative abundance as HCC group predictor. The OOB error in the prediction was 50%. However, the inspection of the top OTU’s that predicted better any differences in the groups, confirmed the identification of the same that in the ANCOM analysis (Figure 5B). Then we selected this three OTU’s as features and trained a new model. This resulted in a classifier with an improved performance, which the testing classification error decreased to 22%.

### 4.6 Sensitivity analysis

We further performed a sensitivity analysis excluding patients under rifaximin (n=8) and lactulose (n=8). We first performed the analysis excluding patients under rifaximin, then excluding patients under lactulose and finally, excluding patients under both medications (n=6). We found that there were not any significant differences related with microbiome profile among cases and controls after exclusion of this important confounder factors (Supplementary Figure 2 A-C and Supplementary Table 1C).

## 5. Discussion

To the best of our knowledge, this is the first observational study to determine the gut microbiome profile among cirrhotic patients with and without HCC. Patients with HCC showed a more diverse gut microbiome than the wo-HCC group. Patients with HCC had specific changes in family members of Firmicutes including a 3-fold increased of Erysipelotrichaceae and a 5-fold decease in family Leuconostocaceae when compared to controls. Second, genus Fusobacterium was found 5-fold decreased in HCC versus wo-HCC. Third, the ratio bacteriodes/prevotella was increased in HCC cases due to the significant decrease in the genus prevotella. This pattern has been associated with an inflammatory milieu with increased activation of NLR signalling pathways.

Microbiome diversity is just one factor to consider when analyzing any ecosystem along with its stability, structure and function. There is currently limited understanding of how informative such measures may be in assessing the state of the gut microbial community. Considering diversity and microbial competition, numerous factors may interact and influence the microbiome stability. In fact, there are some published papers showing that the microbiome is still stable over time and might change in some clinical situations such as decompensated cirrhosis ^2,5^. Many studies have use diversity as a key measure in their analysis, often assuming that diversity correlates with a “healthy” one. This issue might not be always the case. We observed that HCC patients had a less diversity index when compared to healthy controls and a more diverse index when compared to cirrhotic patients without HCC. This might underline the importance of a more complex microbial ecosystem including both, species richness (number of different species in a community) and evenness (relative abundance). We further performed the Tail statistic, which has been specifically developed for 16S rRNA sequence data and has been proved more effective in capturing the diversity among low-abundance species when compared to traditional diversity indices.

Previous observational and small studies, as the present one, have found some changes in the gut microbiome of cirrhosis including a decreased abundance of Lachnospiraceae, Ruminococcaceae and Clostridialies and a higher abundance of Enterobacteriaceae and Bacteriodaceae ^2^. Infection or inflammation triggers changes in gut microbiome with a relative increase in Enterobacteriaceae and Bacteriodaceae ^2,5^. In our study, a higher abundance of Enterobacteriaceae was observed in all cirrhotic patients. Other recent studies have shown that pharmacological intervention can change the type of microbiome and moreover, the use of probiotics such as Lactobacillus decreases inflammatory mediators including tumor necrosis factor (TNF-α) and endotoxemia or LPS ^36–39^. However this microbiome profile in cirrhosis might be once stablished unchanged after the eradication of the injurious factor that originated the liver disease as it was shown in patients after HCV eradication ^39^.

Although inflamation and tumorigenesis have been related in HCC and cirrhosis, a link between changes in gut microbiome, inflamation and liver cancer has only been reported in murine models ^12–14^. In this sense, some specific findings in our study of gut microbiome profile between HCC and wo-HCC patients are important to mention. First, the HCC group showed a more diverse gut microbiome than the wo-HCC group but still less diverse than a healthy population control. This observation could be relevant in the context of disease development and progression suggesting an expansion of certain families or genera of microbes in the tumor microenvironment with an inflammatory milieu. These findings suggest a tumor specific expansion of microbiota ^27^.

Second, as previously mentioned, HCC is a pro-inflamatory cancer ^15^. It is interesting to note that some families in the phylum Firmicutes were either significantly elevated or decreased in HCC cases versus controls wo-HCC. We observed a 3-fold increased of Erysipelotrichaceae in HCC patients. This family of firmicutes has been implicated in inflammation related to gastrointestinal diseases and in tumor development such as colorectal cancer ^28^. Besides, the relative abundance of Erysipelotrichi positively correlated with tumor necrosis factor alpha (TNF) levels in a patients with HIV infection ^29,30^. On the other hand, family Leuconostocaceae, a main producer of acetate and lactate, was decreased by 5-fold in HCC cases versus controls wo-HCC, as also observed in ulcerative colitis ^31^. It is important to mention that the two genera of Firmicutes could potentially serve as predictive biomarker of HCC, since in wo-HCC was decreased by 5-fold and in HCC was increased by 3-fold. Moreover, several members of the gut microbiome were also found significantly decreased in HCC cases. For instance, the genus Fusobacterium was found 5-fold decreased in HCC versus wo-HCC. Elevated levels of Fusobacterium were found associated with tumorigenesis process in the gut and a proinflammatory microenvironment ^32^. However, we found decreased levels in the HCC cases showing that a specific micriobiome related cancer might be found in different tumors.

Third, bacteroides and Prevotella are major genus of the Bacteriodetes phylum present in the gut, but they are mutually excluded in the dominance, as they are clearly competitors for the niche. Prevotella was also linked to chronic inflammation processes. In our study, the ratio bacteriodes/prevotella was increased in HCC cases due to the significant decrease in the genus prevotella and no significant changes in the genus bacteroides. In fact, prevotella was decreased by at least 3-fold in HCC cases. This observation is under further investigation by our group.

Consequently, it seems that in cirrhosis, inflammation and tumorigenesis in HCC might be closely related ^12–14^ In this sense, it has been previously observed that in cirrhosis, a “leaky” gut accounting for an impaired intestinal barrier function and bacterial overgrowth leads to development of main clinical events in cirrhosis. These bacterial products induce inflammation through activation of TLR and NLRs in the liver, inducing a inflammatory environment of cancer development ^12,13^. NOD-like receptors are intracellular sensors with central roles in innate and adaptive immunity. Whereas the membrane-bound Toll-like receptors (TLRs) survey the extracellular milieu, NLRs have emerged as key sensors of infection and stress in intracellular compartments ^14^ These molecules act as sensors of microbial fragments inside the human cytosol of infected cells. Once activated, NLRs recruit NF-κβ and facilitate the production of interferon type I. These NLRs can regulate several aspects of the immune response. In our study, changes in abundance in HCC patients correlated with an increase in the predicted metabolism of NOD-like receptor (NLRs) signaling pathway. However, we are aware that abundant molecular functions are not necessarily provided by abundant species, highlighting the importance of a functional analysis for a community understanding ^14^.

The key question is whether these findings would translate into a clinical relevant knowledge in the field of HCC pathogenesis as a co-carcinogenic factor, and if that is the case, if this changing microbiota profile would be a potentially preventive target for HCC development or recurrence. We tried to validate our findings in a random subset of our HCC/non-HCC patients. We performed a machine learning analysis by the construction of random forest models in order to predict any group from differences in abundance species. Three different OTU’s, 2 from the Odoribacteraceae family, genus Odoribacter and Butyricimonas were identified to be more differentialy abundant in HCC samples. Interestingly, the genus Odoribacter has been linked and associated with colon cancer tumorigenesis ^33^. This finding should be cautiously interpreted because we included different clinical HCC BCLC stages. In this sense, in order to validate our finding we should include patients with very early or early BCLC stages in order to propose these findings as a new biomarker of HCC development. Our group is in a second phase of research including a longitudinal microbiome measurement. This will allow us to clarify whether these preliminary observations explain a novel pathophysiological mechanism in the generation of HCC.

Another sensitivity analysis excluding patients under lactulose or rifaximin was performed, as these drugs could have been potential confounder factors. There were no significant differences in the proportion of cases vs controls under these drugs and in the microbiome profile between cases and controls (without HCC) after excluding patients with lactulose and rifaximine.

Finally, we know that consumption of dietary fibre significantly alters the composition of the intestinal microbiota particularly its diversity. These changes have been associated with a more beneficial taxa (higher Firmicutes and a greater Prevotella:Bacteroides ratio) producing short-chain fatty acids as acetate or butyrate ^35^. However, while long-term diet intervention should be recommended, short-term dietary intervention studies showed rapid but less permanent changes in the microbiome composition with higher inter-subject variability. Furthermore, the impact of dietary habits on colon cancer development initially related with a potentially preventive intervention, opposite results have been recently published showing that microbial-derived butyrate may in fact promote colon polyps development, acting as an oncometabolite ^40^. Thus, while there are several data that supports the changing benefitial microbiome by low-high-fat diets rich in fibre, there is conflicting data regarding cancer development with these interventions. Nevertheless, we consider that randomized clinical trials (RCT) should be mandatory during the next years with longitudinal microbiome assessement.

We recognize that this study has limitations including first, that in case-control studies, although a strict revision of potentially selection and information bias was assessed, several unknown or uncontrolled confounders factors might have influenced the results. Second, the limited number of included patients is a main caveat. Taking this in mind, these preliminary observations might be a starting point to develop a longitudinal assessment of gut microbiota in cirrhotic patients with higher risk of HCC development and prospectively evaluate whether this type of microbiome could be identified as a new HCC biomarker. However, these findings might not be generalizable to all the patients with HCC, especially given the fact that there might have been other factors supporting inter-group and within-group variability. Particularly, specific potential confounder that was not address in our research such as diet ^35^. Although we tried to include a homogenous population with similar dietary habits and precluding high fat diets and alcohol consumption in the eligibility criteria, there was no immediately previous evaluation by any dietician. Nevertheless, although this affects our ability to interpret the data in context of pathophysiology, it bolsters the case for the gut microbiome providing a biomarker of disease. The fact that we were still able to find 3 OTUs that were significantly different in this small cohort without controlling for diet is interesting and remarkable. Our results might be in line with previous observations in other researches regarding the effect of microbiota in cancer development or growth, thus opening a new debate whether the “gut-liver axis” and its inflammatory pathways promotes liver cancer development ^41,42^.

In conclusion, we found several different genera and families in the gut microbiota that were decreased or elevated in HCC cases versus controls (wo-HCC). Patients with HCC had a 3-fold increased of Erysipelotrichaceae and a 5-fold decease in Leuconostocaceae family, a 5-fold decreased abundance in genus Fusobacterium and an increased ratio bacteriodes/prevotella when compared to cirrhotic patients without HCC. This pattern was associated with inflammatory pathways. These findings should be further explored as a novel risk factor for HCC development in a larger cohort study and eventually, in an intervention prevention trial.

## Acknowledgments

Federico Piñero, Patricia Baré and Marcelo Silva received a Clinical Research Grant from the National Institute of Cancer (INC), Argentina.

## Authorship Statement

i. **Guarantor of the article**: Federico Piñero.
ii. **Specific author contributions**: Concept and design, statistical analysis, writing of article: F Piñero, C Rohr, M Vazquez, P Bare, M Silva. Data recording, critical review of the manuscript: M Vazquez, P Bare, M Mendizabal, M Sciara, C Alonso, F Fay and M Silva.
iii. **A statement indicating that all authors approved the final version of the manuscript**: All authors read and approved the final version of the manuscript.

## SUPPLEMENTARY FIGURES

**Supplementary Figure 1.**
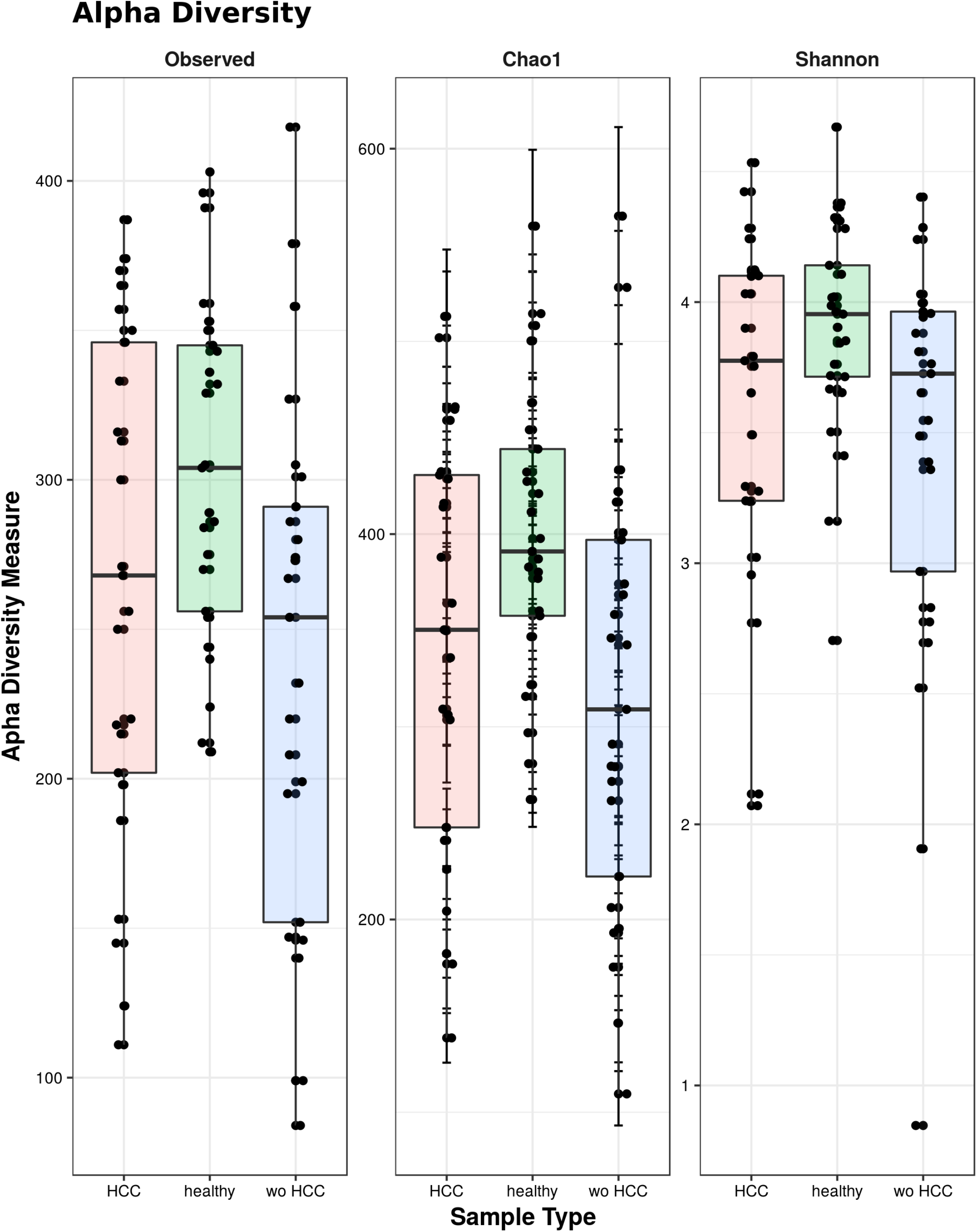
Panel plots comparing alpha-diversity plots between cases (HCC), controls (wo-HCC) and healthy individuals. **Note**: Healthy controls showed the most diverse dataset of gut microbiome. Patients with wo-HCC dataset consistently and significantly showed the less diverse dataset of gut microbiome, while cases (HCC) being less diverse than healthy controls, still more diverse than controls (wo-HCC) dataset.

**Supplementary Figure 2.**
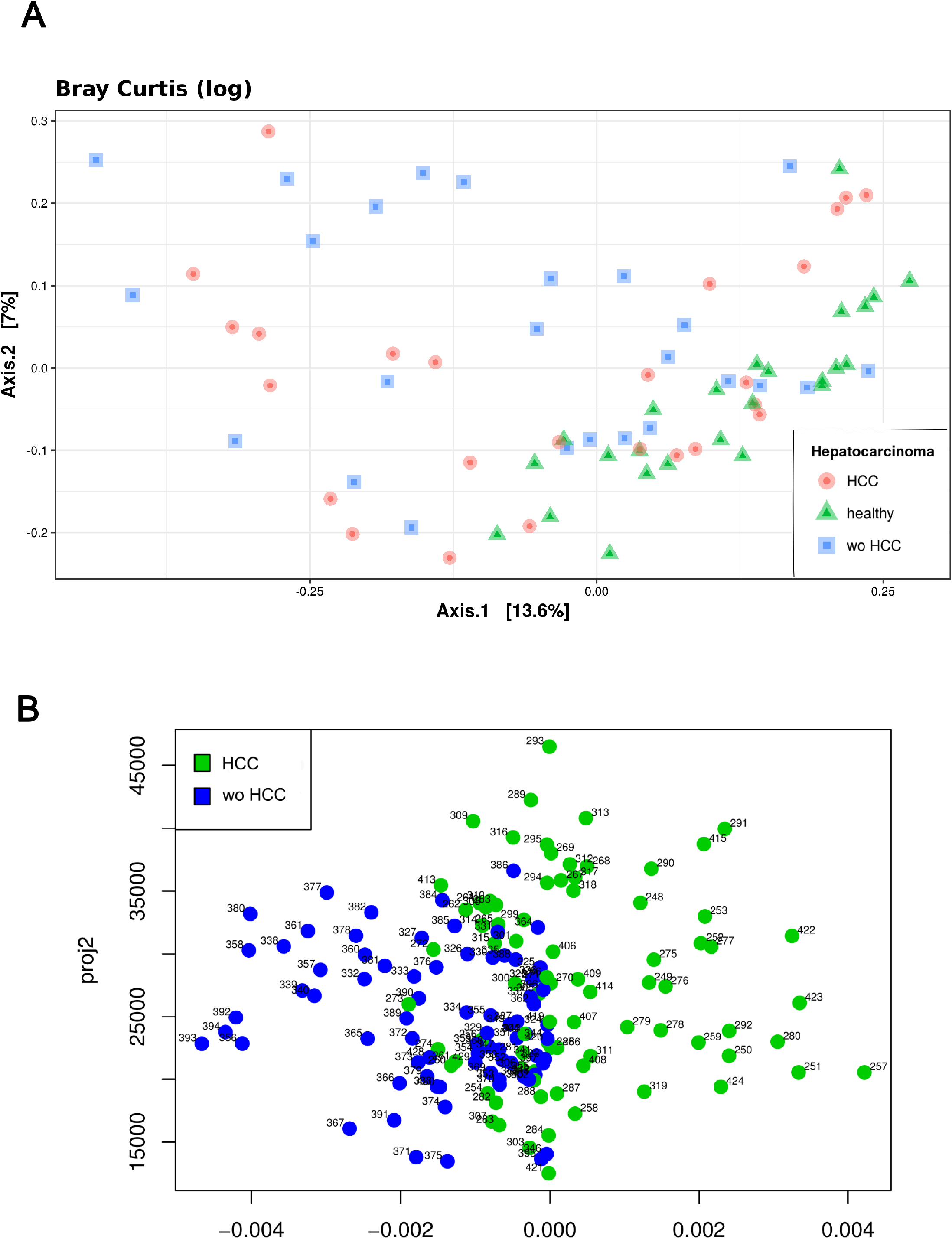
Beta-diversity (Bray Curtis -log; Principal Fitted Components Analysis) plots between cases (HCC), controls (wo-HCC) and healthy individuals. **Note**: Cases and control wo-HCC groups were clearly separated from the healthy control population. A more precise and clear separation of the gut microbiome datasets between groups is shown on the panel below.

**Supplementary Figure 3:**
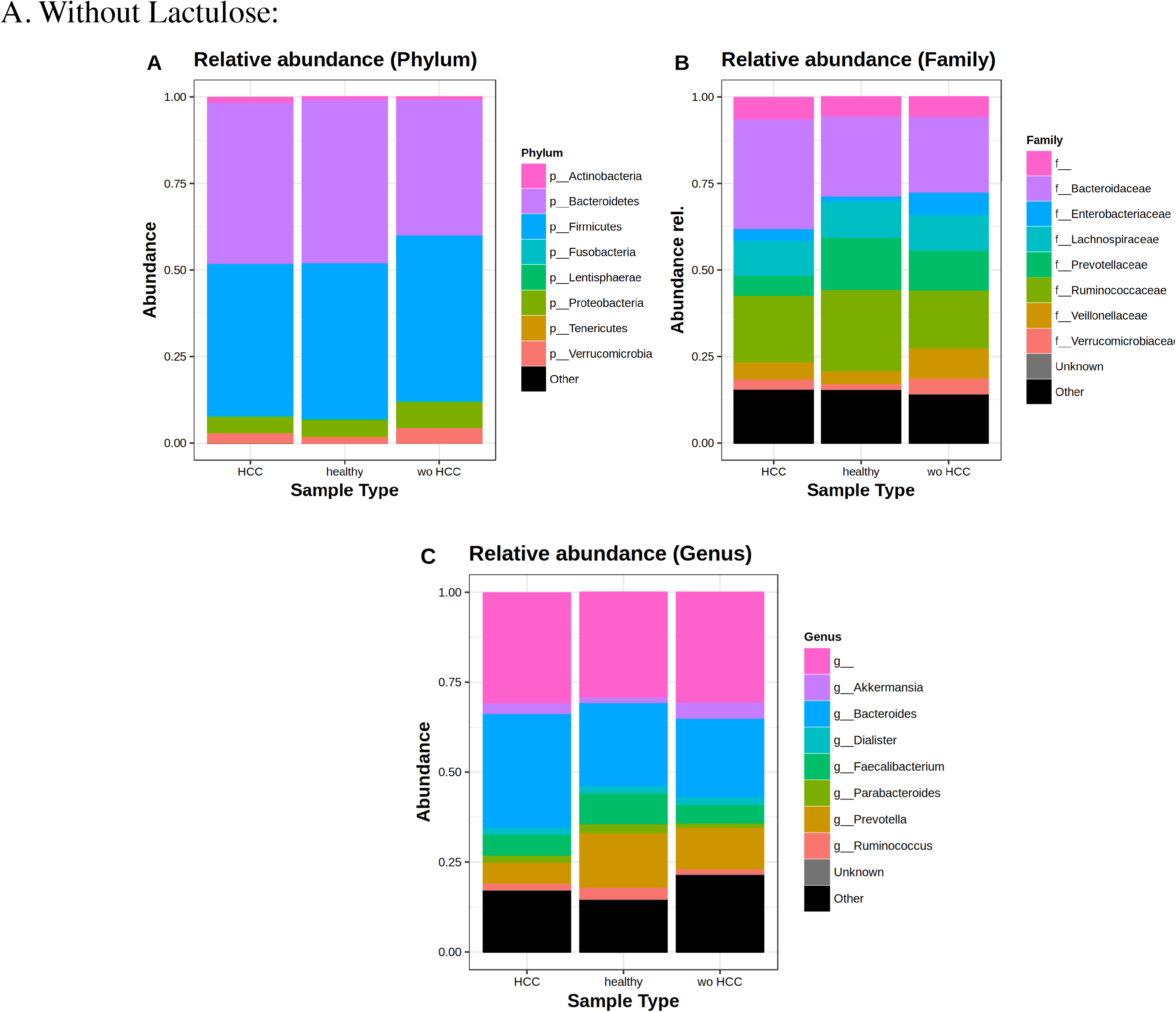
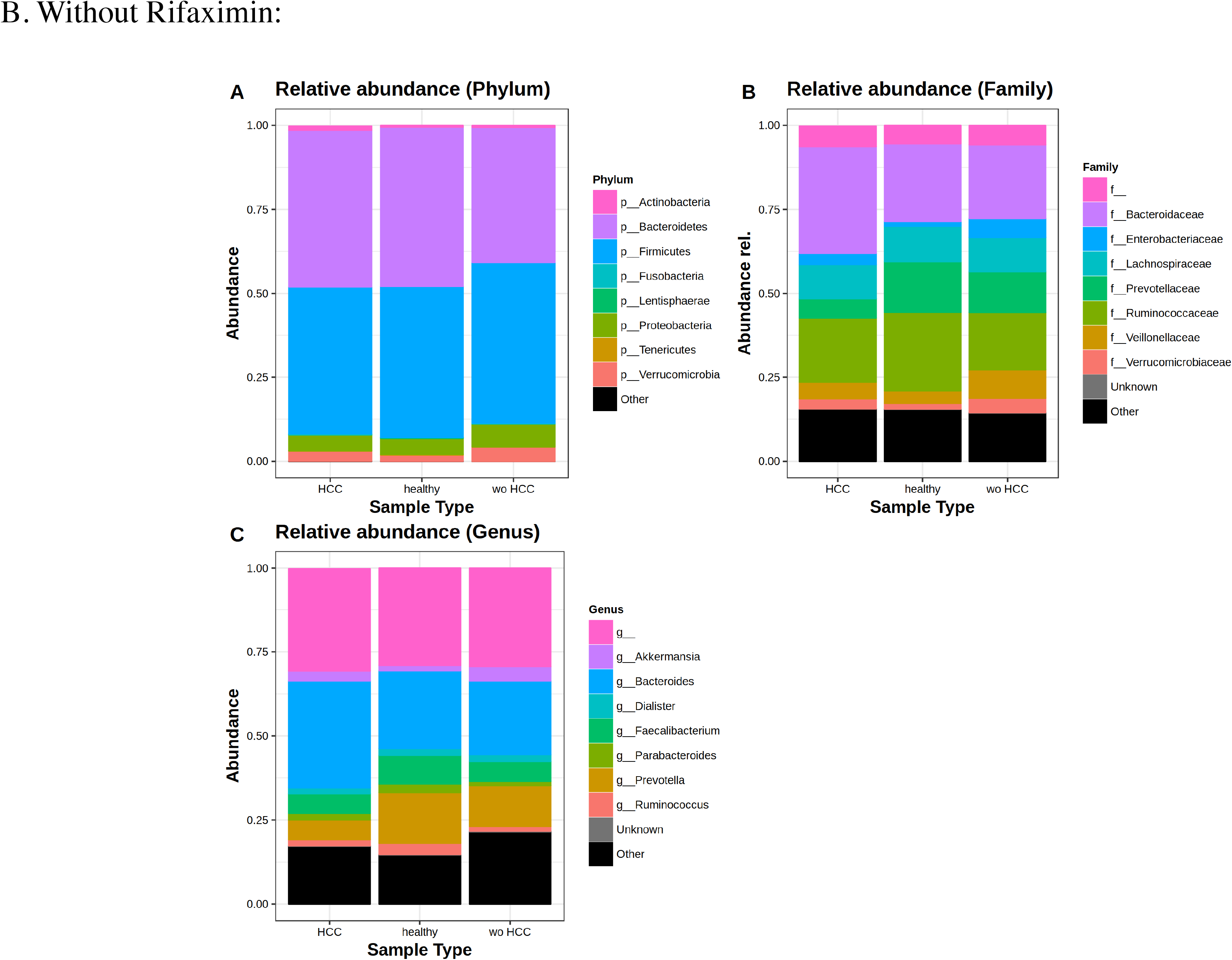
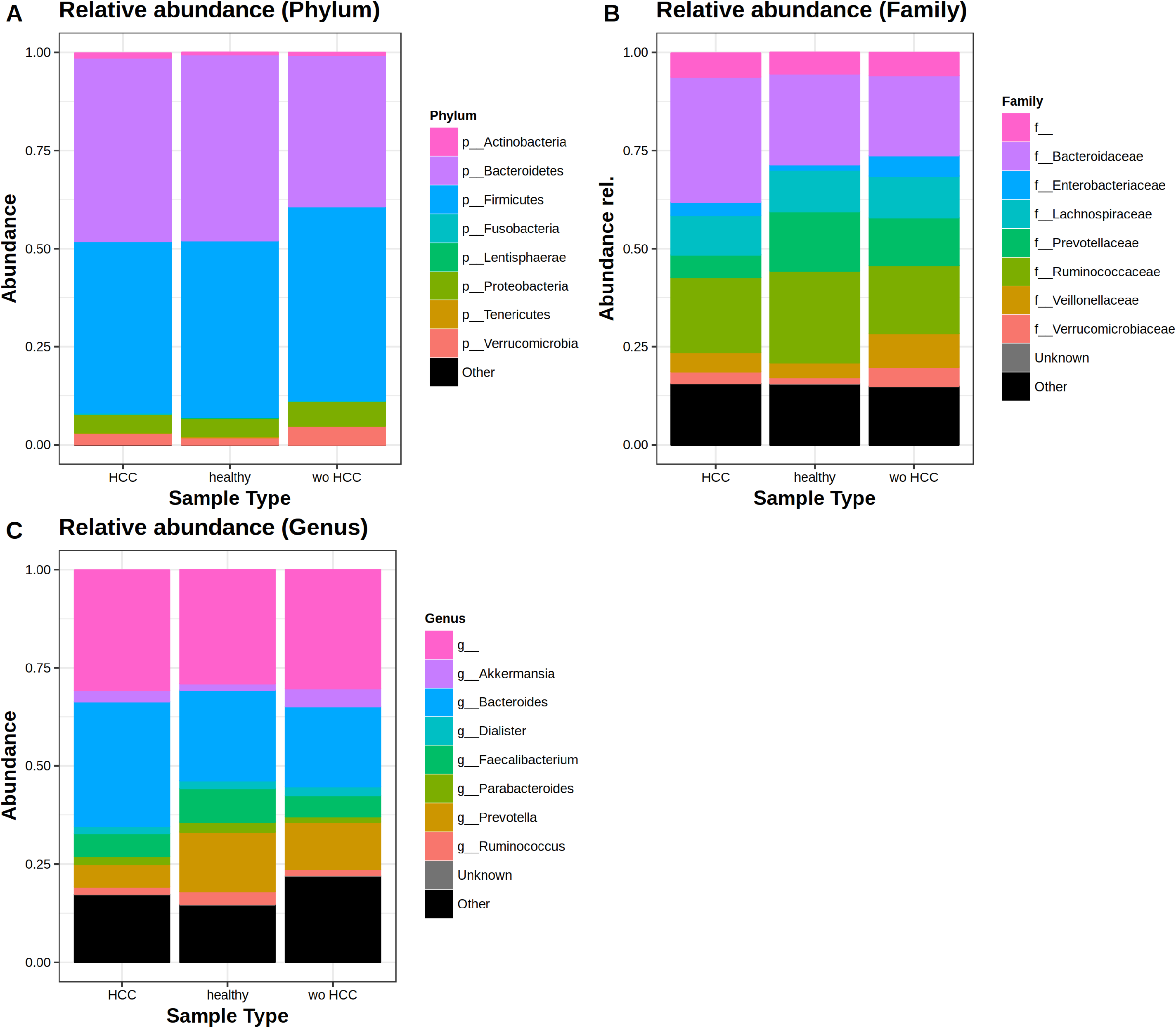
Taxonomic abundance at Phylum, Family and Genus levels between HCC, non-HCC patients and healthy individuals after exclusion of patients under lactulose (Panels A), rifaximin (B) and both drugs (C). A sensitivity analysis excluding patients under lactulose or rifaximin was performed. There were no significant differences in the microbiome profile between cases and controls (without HCC) after excluding patients with lactulose and rifaximine.

**Supplementary Table 1.** A. ANOVA Shannon diversity index healthy vs HCC vs non-HCC.

| **VARIABLE** | *Sum Sq* | *Mean Sq* | *F value* | *Pr (>F)* |
| --- | --- | --- | --- | --- |
| **Hepatocarcinoma** | 2.616 | 1.3081 | 3.068 | 0.04 |
| **Residuals** | 30.702 | 0.4264 |  |  |

**B. Results of Tukey tests between HCC, wo HCC and healthy individuals according to Shannon diversity.**
| **VARIABLE** | *diff* | *95% CI* | *Adjusted P value* |
| --- | --- | --- | --- |
| **Healthy vs HCC** | 0.272 | -0.169;0.714 | 0.31 |
| **Non-HCC vs HCC** | -0.182 | -0.624;0.26 | 0.58 |
| **Non-HCC vs healthy** | -0.454 | -0.896;-0.01 | 0.04 |

**C. ANOVA tests for differences in the Shannon diversity among HCC vs wo HCC groups.**
| **VARIABLE** | *Sum Sq* | *Mean Sq* | *F value* | *Pr (>F)* |
| --- | --- | --- | --- | --- |
| **Hepatocarcinoma** | 0.414 | 0.414 | 0.738 | 0.395 |
| **Male gender** | 0.466 | 0.4658 | 0.83 | 0.367 |
| **Cirrhosis** | 0.878 | 0.8783 | 1.565 | 0.217 |
| **Rifaximine/Norfloxacin** | 0 | 0 | 0 | 0.998 |
| **Lactulose** | 0.186 | 0.1858 | 0.331 | 0.568 |
| **Residuals** | 24.689 | 0.5611 |  |  |

**Supplementary Table 2.** ADONIS on weighted and unweighted UniFrac distances among A) HCC vs wo HCC B) healthy vs HCC vs wo HCC.

| *Unweighted UniFrac* |  |  |  |  |  |
| --- | --- | --- | --- | --- | --- |
| **VARIABLE** | *Sum Sq* | *Mean Sq* | *F model* | *R2* | *Pr (>F)* |
| **Hepatocarcinoma** | 0.194 | 0.1943 | 1.1792 | 0.0241 | 0.218 |
| **Male gender** | 0.133 | 0.1329 | 0.8068 | 0.0165 | 0.721 |
| **Cirrhosis** | 0.152 | 0.1522 | 0.824 | 0.0188 | 0.528 |
| **Residuals** | 7.58 | 0.1647 |  | 0.94 |  |
| *Weighted UniFrac* |  |  |  |  |  |
| **VARIABLE** | *Sum Sq* | *Mean Sq* | *F model* | *R2* | *Pr (>F)* |
| **Hepatocarcinoma** | 0,0367 | 0,0367 | 0,9069 | 0,0188 | 0,488 |
| **Male gender** | 0,0284 | 0,0284 | 0,7026 | 0,0145 | 0,68 |
| **Cirrhosis** | 0,0238 | 0,0238 | 0,5886 | 0,0122 | 0,812 |
| **Residuals** | 1,8652 | 0,04 |  | 0,9543 |  |

**B.**
| *Unweighted UniFrac* |  |  |  |  |  |
| --- | --- | --- | --- | --- | --- |
| **VARIABLE** | *Sum Sq* | *Mean Sq* | *F model* | *R2* | *Pr (>F)* |
| **Hepatocarcinoma** | 0,8135 | 0,4067 | 2,7372 | 0,0706 | **0,001\*** |
| **Residuals** | 10,6989 | 0,1486 |  | 0,9293 |  |
| *Weighted UniFrac* |  |  |  |  |  |
| **VARIABLE** | *Sum Sq* | *Mean Sq* | *F model* | *R2* | *Pr (>F)* |
| **Hepatocarcinoma** | 0,1354 | 0,0677 | 2,0334 | 0,0534 | **0,017\*** |
| **Residuals** | 2,3983 | 0,0333 |  | 0,9465 |  |

## Abbreviations

AFP: Alpha-fetoprotein
BMI: Body Mass Index
CC: cryptogenic cirrhosis
CI: Confidence Interval
CT: Computerized Tomography
HCC: Hepatocellular carcinoma
HR: Hazard ratio
IQR: :interquartile range
LT: Liver Transplantation
MC: Milan criteria
MELD: Model for End-stage Liver Disease
MRI: Magnetic Resonance Imaging
NAFL: Non-alcoholic fatty liver
NAFLD: non-alcoholic fatty liver disease
NASH: Non-alcoholic steatohepatitis
NLR: NOD-like receptors
OTUs: Operational Taxonomic Units
PEI: Percutaneous ethanol injection
RFA: radiofrequency ablation
TACE: trans-arterial chemoembolization
TLR: Toll-like receptors

